# Computed Tomography of Polymeric Biomedical Implants from Bench to Bedside

**DOI:** 10.1101/2024.02.20.581229

**Authors:** Kendell M Pawelec, Todd A Schoborg, Erik M Shapiro

## Abstract

Implanted biomedical devices require porosity to encourage tissue regeneration. However, characterizing porosity, which affects many functional device properties, is non-trivial. Computed tomography (CT) is a quick, versatile, and non-destructive way to gain 3D structural information. While optimization of CT for polymeric devices has been investigated at the bench on high-resolution micro-CT (μCT) scanners, pre-clinical and clinical systems cannot be tuned the same way, given an overriding objective to minimize ionizing radiation exposure to living tissues. Therefore, in this study we tested feasibility of obtaining structural information in pre-clinical systems and μCT under physiological conditions. The size of resolved features in porous structures is highly dependent on the resolution (voxel size) of the scan. Lower resolution underestimated porosity and overestimated pore size. With the homogeneous introduction of radiopaque nanoparticle contrast agent into both biopolymers and synthetic polymers, devices could be imaged in the hydrated state, even at high-resolution. Biopolymers had significant structural changes at the micro-scale post-hydration, including a mean increase of 130% in pore wall thickness that could potentially impact biological response. Through optimizing devices for medical imaging, CT has the potential to be a facile way to monitor devices from initial design stages through to clinical translation.

## 1. INTRODUCTION

A fundamental design criterion for implanted tissue engineering devices is the ability to stimulate the patient’s natural regenerative capacity to repair functional tissue after trauma. This requires engineering of devices that can support the infiltration of native cells and vasculature at the injury site and therefore drives the use of porous materials. While the need for porous materials cannot be denied, the introduction of porosity also affects biomedical device mechanics [1] and can introduce anisotropy in materials properties like permeability [2]. Porosity is a key feature of devices that must be characterized. However, the measurement of 3D features can be difficult, particularly as devices increase in complexity. Thus, computed tomography (CT) has become a useful tool for engineers and clinicians for its non-destructive nature, versatility, speed, and high spatial resolution.

CT relies on the ability of materials to interact with, or attenuate, x-rays, to create 3D images, differentiating materials and tissues based on differences in attenuation coefficients [3]. At the highest resolutions, micro-CT (μCT) has a proven reproducibility and accuracy compared to destructive techniques, such as confocal scanning laser microscopy, when assessing tissues such as cortical bone [4]. Importantly, CT also has an established track record of use in the clinic, giving it the potential to be a pertinent imaging technique of novel biomedical devices throughout translation from the lab bench to the bedside [5].

In practice, reliable CT data can be complicated by many factors including the sample type, its environment, acquisition parameters and post-processing steps [6,7]. A common problem is the accurate imaging of polymeric materials, which form the basis of many biomedical devices. Their low x-ray attenuation has resulted in numerous optimization studies at high resolution [7,8], but these acquisition steps do not translate well into the imaging of living biological systems. In particular, hydrated polymer devices cannot be distinguished from tissue, necessitating the integration of biocompatible and homogeneous contrast agents into devices to facilitate a cohesive platform of CT imaging from bench to bedside [9].

Here we investigate practical guidelines for how the transition from high resolution μCT to pre-clinical CT affects the type and accuracy of device features that can be quantified. This was accomplished utilizing phantoms mimicking biomedical devices: a freeze-dried biopolymer and a salt-leeched synthetic polyester [10,11]. Further, the use of a biocompatible radiopaque nanoparticle contrast agent, based on tantalum oxide [12], was shown to allow detection of micron-scale features in hydrated environments in both biopolymers and polyesters.

## 2. MATERIALS AND METHODS

The study utilized phantoms made from composites of polymer + 20wt% nanoparticle contrast agents, designed to mimic biomedical devices. The polymer matrix was either a biological polymer (collagen) or synthetic polymer (polyester).

### 2.1 Radiopaque contrast agent

The contrast agent used to confer radiopacity to composite structures was a tantalum oxide (TaOx) nanoparticle. Nanoparticles were formed by a micro-emulsion technique, as described previously [12]. Hydrophobicity of the particles was tuned via the selective addition of commercially available silanes. Hydrophilic nanoparticles were formed with the addition of 2-[methoxy(polyehyleneoxy)-9-12-propyl]trimethoxysilane (PEG-silane), followed by (3-aminopropyl)trimethoxysilane (APTMS), and subsequent surface modification using methoxy-poly(ethylene glycol)-succinimidyl glutarate (m-PEG-SG-200) in ethanol. Hydrophobic TaOx nanoparticles received a surface coating of PEG-silane and hexadecyltriethoxysilane (HDTES, Gelest Inc., cat no SIH5922.0).

### 2.2 Synthetic polymer composite

Polyester phantoms were created with micro-scale porosity (< 90μm) and macro-scale porosity (> 90 μm) to mimic tissue engineering constructs which must accommodate both nutrient diffusion and cell and tissue infiltration [13]. The synthetic polymer matrix chosen was polycaprolactone (PCL, Sigma Aldrich), a commonly used biocompatible polyester [14]. PCL was solubilized in a suspension of TaOx nanoparticles to create an 8wt% solution in dichloromethane (DCM, Sigma). The nanoparticle suspension was calculated to be 20wt% TaOx of the final phantom mass (polymer + nanoparticles). Sucrose (particle size 31 ± 30 μm, Meijer) was added to the suspension, at 70 vol% of the polymer + nanoparticle mass in solution, followed by sodium chloride (NaCl, Jade Scientific) at 60 vol% of the total polymer + nanoparticle volume. The suspension was vortexed and pressed into a silicon mold (4.7 mm diameter, 2 mm high). After air drying, phantoms were trimmed of excess polymer composite and washed for 2 hours in distilled water, changing the water every 30 minutes to remove sucrose and NaCl. Washed phantoms were air dried and stored in a desiccator prior to imaging.

Microporous films were created from PCL, solubilized at 4wt%, in a suspension containing TaOx nanoparticles. The amount of TaOx was calculated to be 20wt% of the total dry mass (polymer + nanoparticle). Sucrose was added to the solution (70 vol%) and suspensions were vortexed and cast on glass sheets. Films were dried at room temperature, removed and submerged in Milli-Q water to remove the sucrose, then dried for storage.

### 2.3 Biopolymer composite

The biopolymer chosen for radiopaque phantoms was collagen from bovine Achilles tendon (C9879, Sigma), produced via a lyophilization method reported previously [15,16]. Collagen (2wt%) was suspended in a 0.05 M acetic acid solution containing 20wt% hydrophilic TaOx nanoparticles overnight at 4°C. After, a slurry was prepared by homogenizing the solution for 30 minutes (4000 rpm) on ice. Suspension was pipetted into wells of a 24 well tissue culture plate (0.75 ml each), and stored at 4°C until freezing in a Virtis freeze drier at -30°C for 90 minutes, with a cooling rate of 0.9°C/min. Biopolymer phantoms were lyophilized at 0°C for 20 hours under a vacuum of < 100 mTorr, and stored at room temperature prior to imaging.

### 2.4 Radiological Model of Implantation

Two radiological models of implanted biomedical devices were used that contained physiologically relevant tissue structures (bone, fat, muscle): a chicken breast and a food-quality lamb shank. In preparation, polyester films were cut to 5 mm × 20 mm and hydrated overnight in Milli-Q water. The hydrated films were wrapped around a 6 mm agarose gel cylinder and adhered with surgical tape. The gel was then removed and a prepared biopsy of tissue was inserted. The assembly was sonicated briefly in water to remove air prior to implantation. A slit was made in the tissue, and a 6 mm biopsy of tissue was removed at the bottom of the slit. The prepared film was inserted into the cavity and flooded with phosphate buffered saline (PBS), before securing the implant and suturing the slit closed with vicryl (braided, 3-0, FS-1 reverse cutting, J452H Ethicon). The tissue models were stored at 4°C until imaging. Chicken breasts with implants were imaged with a pre-clinical μCT, and lamb shanks were imaged on a clinical CT system.

### 2.5 Scanning electron microscopy

Samples were adhered to 13mm aluminum stubs and sputter coated with gold (≈ 30 nm thickness) in an Emscope Sputter Coater model SC 500 (Ashford, Kent, England) purged with argon gas. Samples were examined in a JEOL 6610LV (tungsten hairpin emitter) scanning electron microscope (JEOL Ltd., Tokyo, Japan), at 12kV and a working distance of 12mm.

### 2.6 Computed Tomography (CT)

#### High-resolution Micro Computed Tomography (μCT)

Dry phantoms were punched to 1.5 or 3 mm diameter. Imaging was performed using a ZEISS Xradia 610 Versa High-Resolution 3D X-ray Tomography Microscope. Samples were imaged in P20 pipette tips in both the dry and hydrated state (1x phosphate buffered saline) at 80 keV, 125 μA (10W) with a 1.2-4.7 mm field of view corresponding to 1.2-4.7 μm resolution using the 4x objective. 1601 or 4501 projection images were acquired in 360-degree scans, with 0.25-1.5 second camera exposure times depending on the sample. Scan times ranged from 38 minutes to 4 hours max. Tomograms were generated using Scout-and-Scan™ Control System Reconstructor Software.

#### Pre-clinical μCT

All tomography images were obtained using a Perkin-Elmer Quantum GX. In vitro phantoms were imaged in microcentrifuge tubes, at 90 keV, 88 μA, with a 36 mm field of view at a 50 μm resolution. They were imaged both in a dry state and after hydration in phosphate buffered saline (PBS). After acquisition, individual phantoms were sub-reconstructed using the Quantum GX software to 12 μm resolution. For standard resolution scans, a 2 min scan time was used (step size: 0.11 degrees, exposure time per projection: 36 msec); high-resolution scans utilized a 57 min scan time (step size: 0.01 degrees, exposure time per projection: 133 msec). Phantoms implanted in chicken breast were scanned at: 90 keV, 88 μA, with a 72 mm field of view, at 90 μm resolution.

#### Clinical CT

Imaging was performed on a clinical CT (GE Discovery, 64 slice). Scans of the entire lamb shank were taken at 120keV, following the standard clinical protocol for a head scan. The resulting images had a resolution of 430 μm/voxel.

### 2.7 Tomography image analysis

Tomography images were acquired from 3-5 independent phantoms per group. The polymer matrix was segmented from background using a histogram of intensities. From the tomography scans of phantoms, several measures of scaffold structure and porosity were quantified. Pore size and overall percentage porosity of the phantoms was analyzed using Image J. Image stacks were opened and rotated, using built in functions, to perform calculations on planes perpendicular to the transverse cross-section in 4-10 slices throughout the phantom, which were averaged for the mean value per phantom. Prior to using built in particle size analysis tools, stacks were converted to binary images by thresholding, the image was despeckled, and a watershed operation was applied to close off adjacent pores. For synthetic polyester phantoms, pores < 90 μm were termed “micro-porosity” and all others were “macro-porosity”. The level of 90 μm was chosen after analysis of a microporous polyester film, showing that 99% of all pores were < 90 μm within the material. Analysis of the polymer matrix thickness before and after hydration was performed using Dragonfly (v.2022.2, Comet Technologies Canada Inc.) built in functionality. The mean of the thickness value computed from the histogram is reported.

### 2.8. Statistics

Statistics were performed using GraphPad Prism (v. 10.0.2). All data was analyzed via ANOVA, followed by Fisher’s LSD test. In all cases, α < 0.05 was considered significant, with a 95% confidence interval. Data are presented as mean ± standard deviation.

## 3. RESULTS and DISCUSSION

Materials and devices have different properties depending on the length scale that is being examined. The same is true of biomedical devices and tissue structures, which are often hierarchical. Naturally, the greatest possible image resolution is sought, to ensure reliability of feature measurement [6]. However, assessing the finest features comes at a cost: massive files that require high computing power to analyze and large time commitments for data acquisition [7].

### 3.1 Pre-Clinical Tomography Resolution

As biomedical devices transition from pre-clinical small animal models to large animal models or humans, the features that can be resolved and measured in CT scans is impacted, Figure 1. This is chiefly due to the limitations on image resolution as the scanner size increases. Realistically, the features that can be measured in each type of machine are 2-3 times greater than the smallest voxel size [17]. Thus, while a high resolution μCT can resolve even the microporosity of a polymer film, pre-clinical systems can only differentiate film thickness and clinical systems are generally limited to overall device dimensions.

**Figure 1.**
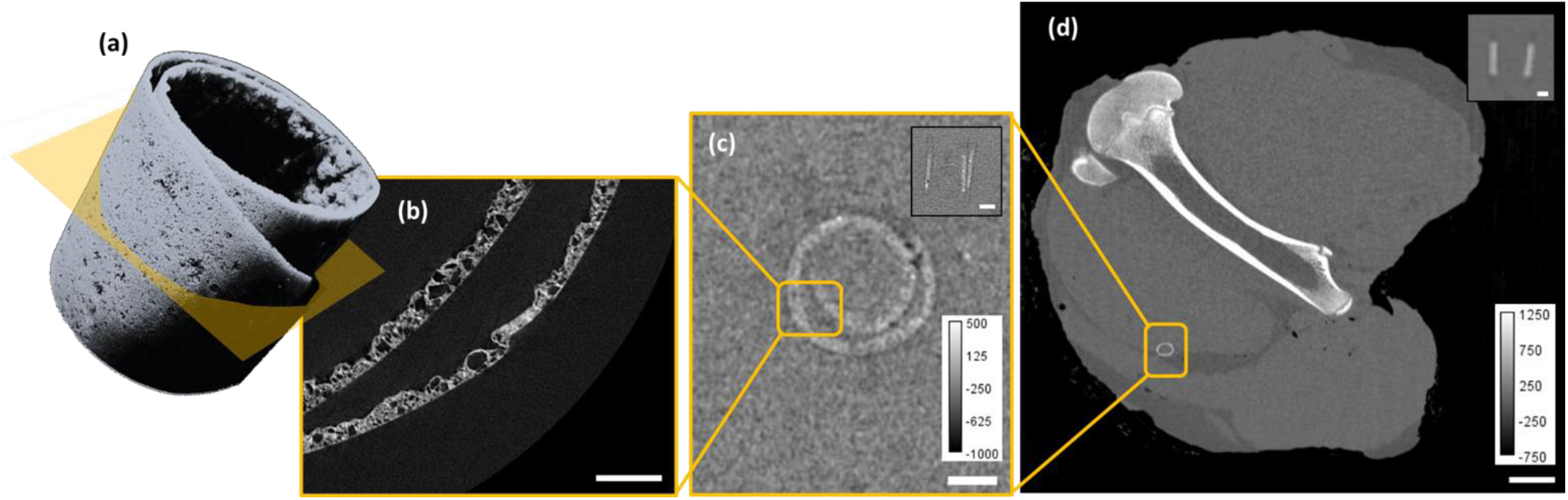
Transitioning from pre-clinical to clinical CT systems brings a decrease in resolution and features that can be resolved. A radiopaque microporous film (180 μm thick) was visualized using tomography and is (a) shown as a 3D reconstruction with the transverse cutting plane marked in orange. The film was hydrated and imaged. Transverse slices are shown from a (b) high resolution μCT (voxel < 5 μm) and after implantation into tissue, using (c) a pre-clinical μCT scanner (voxel size 90 μm) and (d) a clinical CT (voxel size 430 μm); insets show a longitudinal slice. Scale: (b) 250 μm, (c) 2 mm, (d) 20 mm, insets: 2 mm.

The drop in resolution with pre-clinical and clinical CT systems is due to the constraint to minimize exposure to ionizing radiation in living tissue [18]. Thus, when using CT for monitoring living systems, acquisition modes are optimized by CT manufacturers to give the greatest resolution with a minimum radiation exposure per scan [19]. Therefore, many acquisition parameters are preset. Voltage and current are standardized between 80-140 keV in clinical and pre-clinical settings, and step size and exposure time are preset as scan protocols relating to “standard” or “high resolution”. In contrast, benchtop high-resolution μCT scanners have the option of varying a broader range of acquisition parameters: voltage, current, power, exposure time, step size [8]. Indeed, the majority of studies on imaging polymer structures has been optimized to use low voltages (< 50 keV) that are not practical for transition to clinical CT protocols [7].

In porous biomedical devices, two commonly used metrics to describe structure are pore size and porosity, both of which were significantly impacted by the scan resolution in a pre-clinical CT system, Figure 2. Pore size was overestimated at lower resolutions, with no significant changes in measurement with increased scan time, which relates to a decreased step size (0.01 vs 0.1 degrees). The pore size measured was 206.3 ± 16 μm at a resolution of < 5 μm compared to 551.4 ± 39 μm and 514.5 ± 26 μm for 50 μm voxel size at 2 min and 57 minute scan times. The overestimation of pore sizes has been observed in other studies and is related to partial volume effects during segmentation [6,7]. In agreement with the current study, decreases in step size (related to time of scan for the present case) have been shown to play a much less significant role on porous features than voxel size in high-resolution benchtop scanners [7].

**Figure 2.**
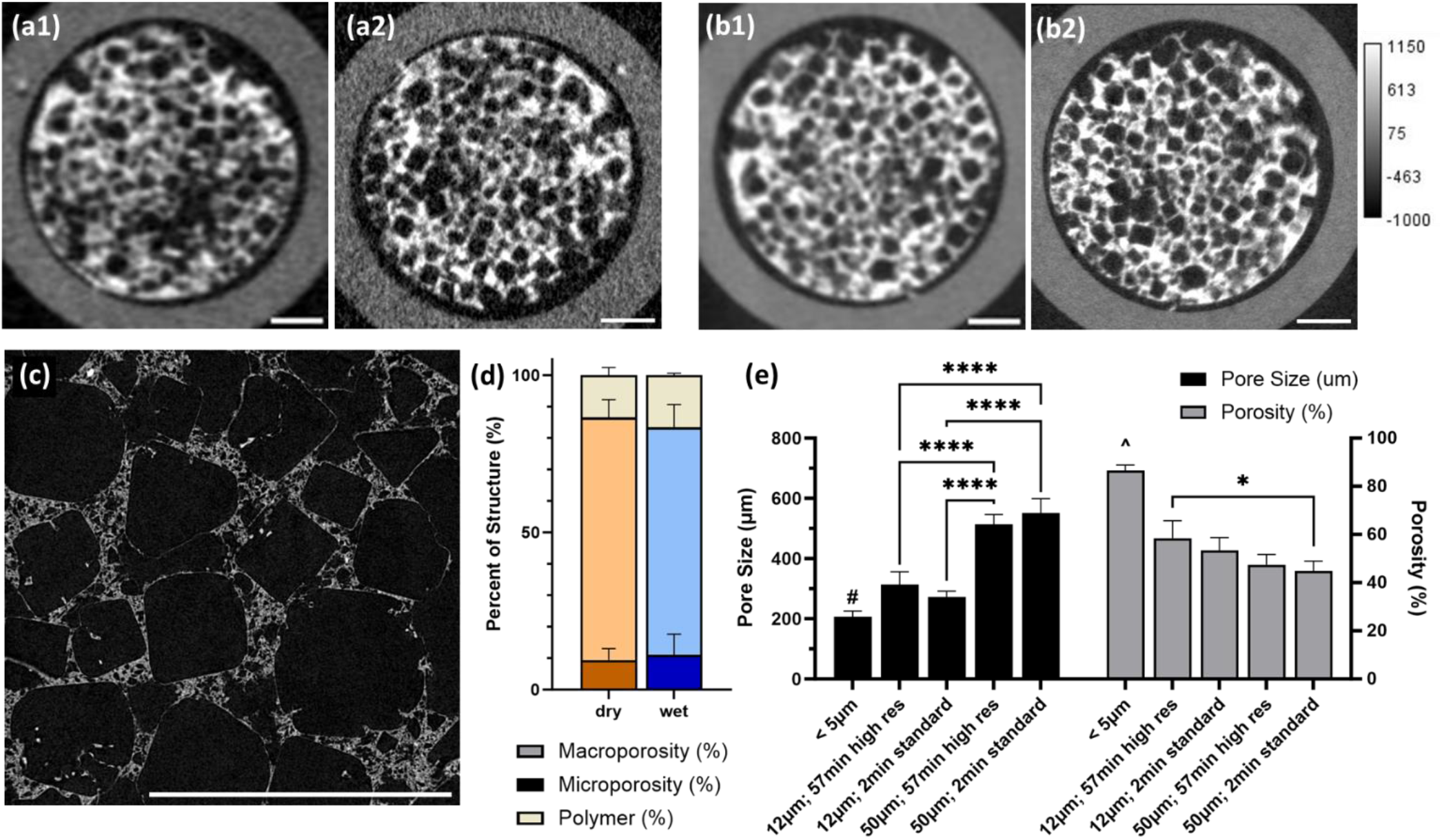
The measured size of porous features in polymer implants is more dependent on the resolution of the μCT scan than the scanning time. Porous polyester phantoms were scanned with (a) a standard 2 min scan (step size: 0.11 degrees) and (b) a 57 min high resolution scan (step size: 0.01 degrees) with a voxel resolution of (1) 50 μm and (2) 12 μm. Only at high resolution (< 5 μm) are both the micro and macroporous features visible and (d) quantifiable in the dry and hydrated state. (e) A larger voxel size led to significant increases in measured pore size and decreases in percent porosity. * p < 0.05, **** p < 0.0001, # significantly different from all other groups (p < 0.001), ^ significantly different from all other groups (p < 0.0001). Scale: (a-c): 1 mm.

Percent porosity was under reported at larger voxel sizes, as the microporous features could not be resolved. In the worst case, 50 μm and 2 min scan time, measured porosity of the phantom was only 44.9 ± 3% of the total volume, compared to a value of 86.5 ± 2% at a voxel size of < 5 μm. Increased scan time did have a significant benefit for measured porosity. However, it was only at 5 μm voxel size that the overall porosity could be segmented into microporosity and macroporosity, Figure 2(d). The macroporosity alone accounts for 77. 1 ± 5% of the total porosity, still significantly greater than the 58.5 ± 6% measured by the next best resolution of 12 μm, 57 min scan time. In bone microstructures, as in the present case, larger voxel sizes are known to significantly underestimate calculated porosity [20]. For rodent bone, even slight differences of 1 vs 2 μm voxel size can significantly alter quantified metrics and the ability to resolve the finest features [20]. Some groups have suggested utilizing equations to adjust the values obtained from lower resolution scans in order to calculate a “correct” value while minimizing file size and scan time, requiring a measure of the under- or over-estimation [7]. Thresholding at various resolutions might also impact the porosity measurements. However, it has been noted that changes to features by alternative thresholding techniques does not play nearly as large a role in variation as voxel size [17].

While calculated feature size may be impacted by scan resolution, the ability to differentiate different biological effects may still remain in low resolution data, even if absolute values vary significantly from true metrics, so long as the standard deviation of measured parameters does not also increase at lower resolution [6]. However, in some instances, such as creating computer models, all features must be distinguished at the highest resolution possible. For example, models of biomedical device mechanics rely on accurate measures of overall porosity as the contributing factor for determining elastic modulus [21], and insufficient resolution could skew model predictions.

Within tissue engineering and biomedical devices, a variety of polymeric materials utilized is extensive. While they have vastly different properties, polymers generally all have poor x-ray attenuation properties. Given the host of different manufacturing techniques that can form porous structures, depending on chemical compatibility and device characteristics, the features of biomedical devices can change dramatically. With this change, so does the information that can be reliably acquired by CT techniques.

One example of the diversity of porous polymer devices can be illustrated with the comparison of structures made from a biopolymer and synthetic polyester, Figure 3. A classic technique for forming porous structures from biopolymers, given their hydrophilic nature, utilizes freeze drying [15]. Implants produced in this way exhibit high interconnected porosity with very fine biopolymer walls in an architecture that is tuned via freezing parameters [16,22]. Even macroscopic features are difficult to resolve in pre-clinical μCT due to the fine wall thickness.

**Figure 3.**
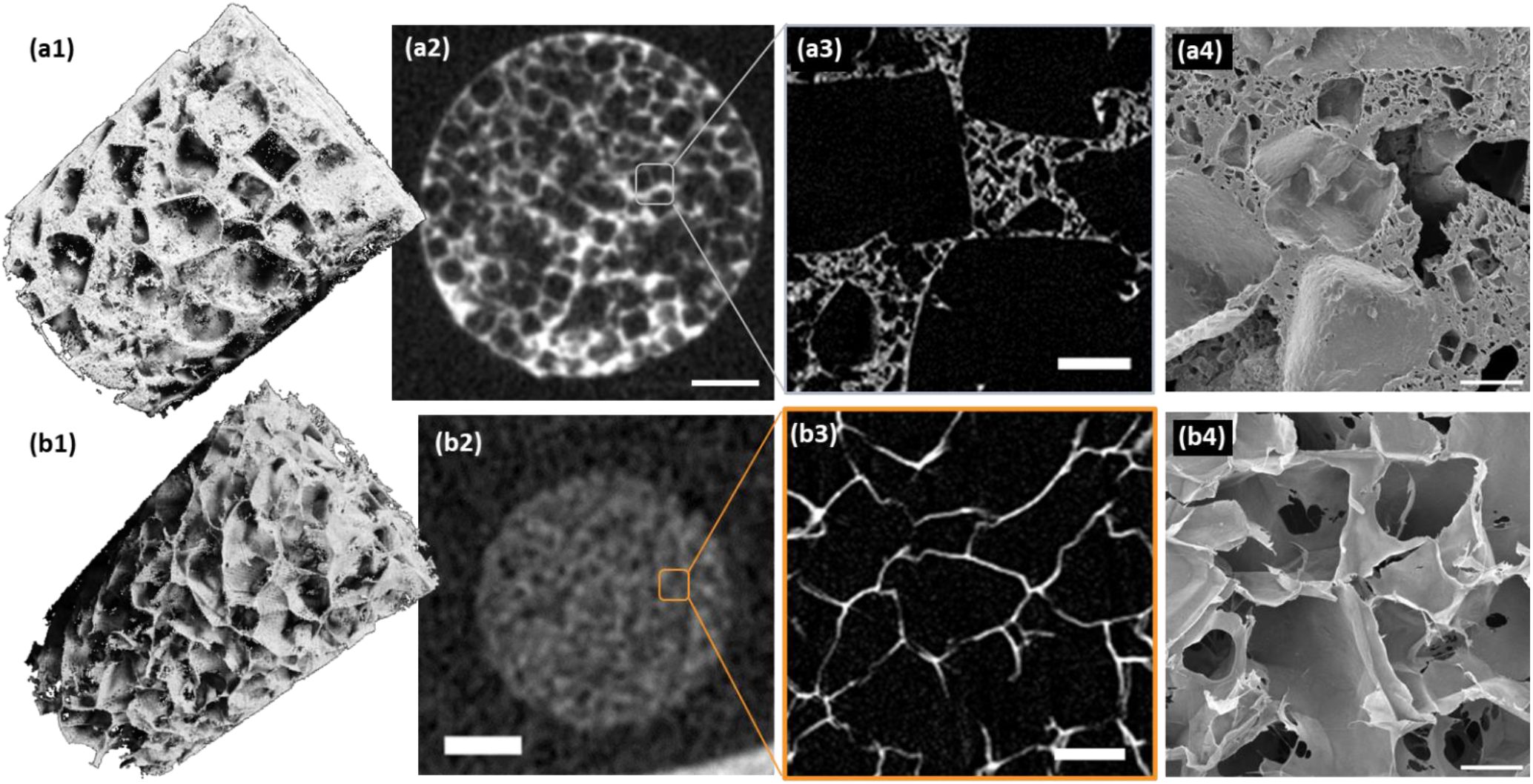
Material type influences manufacturing type and the resolution required to accurately quantify porous features. Porous phantoms were made from (a) a polyester (PCL) and (b) a biopolymer (collagen). The phantoms had distinct features, as observed via (1) a 3D reconstruction of a high resolution μCT scan, (2) 12 μm voxel size and (3) 1.5 μm voxel size. (4) The porous features observed in the μCT were verified by scanning electron microscopy (SEM). Scale: (2) 1 mm, (3) 100 μm, (4) 100 μm.

By comparison, salt leaching or 3D fused filament fabrication (FFF) techniques result in larger wall thicknesses that can be resolved as porous structures even at low scan resolution, Figure 3(a). However, while features are finer, the same trends in feature size estimation that occur in polyester systems have been shown to occur in biopolymer systems as well, as voxel size varies from 1-15um [7].

For any measurement of porosity, validation with other characterization techniques is required. Most features should correlate, but post-processing may contribute to bias in CT data [4]. As a check, macroscopic device properties can be used, for example fluid permeability can support measured values of interconnectivity [2]. Often, validation is done with other forms of destructive microscopy, such as confocal microscopy [23], or scanning electron microscopy (SEM) that can be paired with focused ion beam (FIB) to achieve 10 nm resolution over small volumes [24].

### 3.2 Devices in Hydrated Environments

While a major advantage of CT is its ability to be utilized for samples in a variety of states, subjected to outside forces or undergoing phase transitions, the utilization of μCT to determine porous features of polymer devices is generally restricted to dry systems. The low attenuation of polymers means that in an aqueous environment the attenuation of polymer and water is indistinguishable, Figure 4(a). However, biomedical devices are designed to work in biological systems, and for certain classes of materials, such as biopolymers, hydration has a huge impact on materials properties [25].

**Figure 4.**
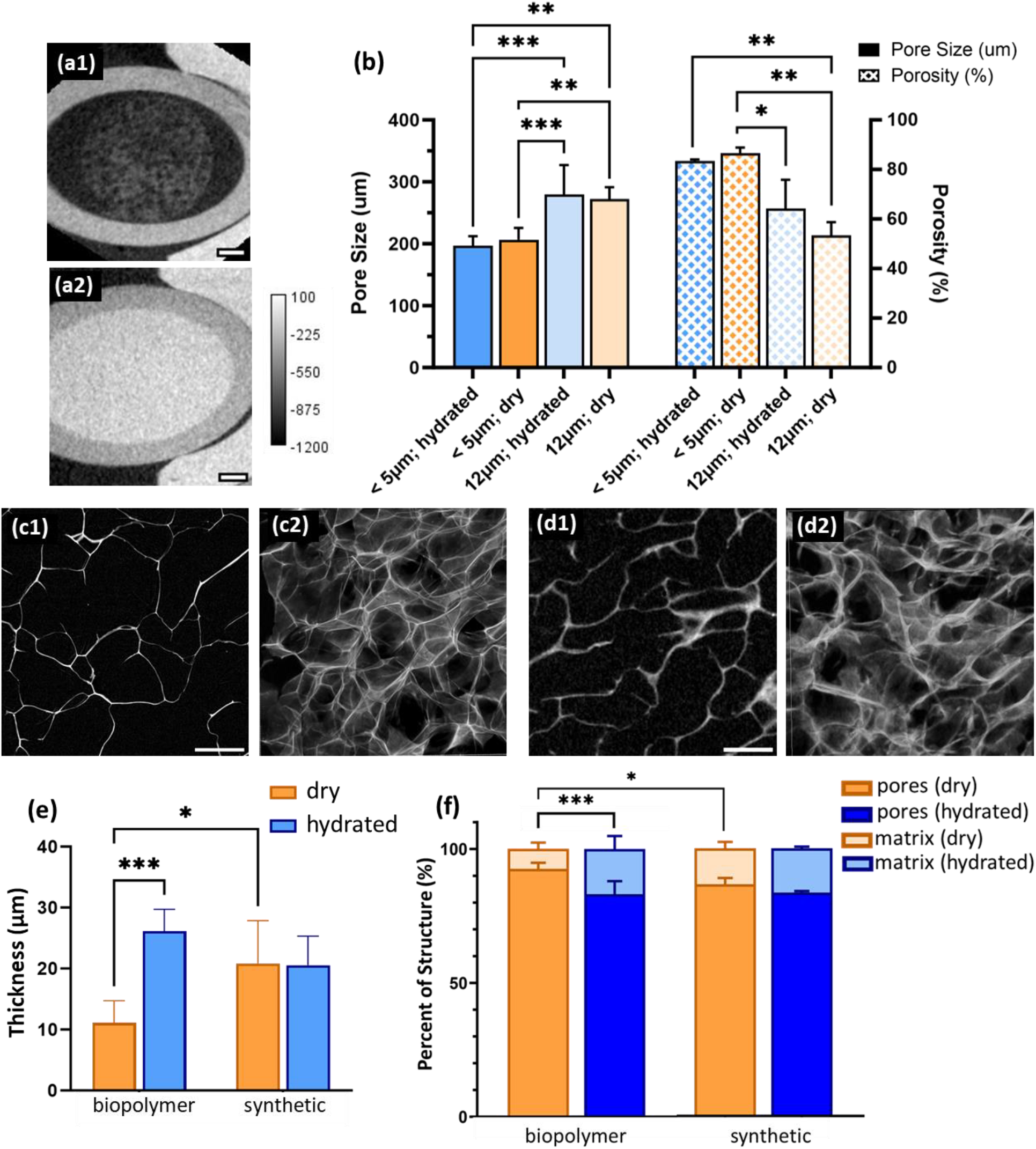
Tomography allows for non-invasive imaging of samples in various environments, such as post-hydration. (a) Imaging polymers without contrast agents in a hydrated state is challenging given lack of contrast with the aqueous environment: (1) dry PCL phantom (0wt% nanoparticles) and (2) hydrated. Homogeneous incorporation of nanoparticle contrast agent allows (b) quantification of features in both states, with features size again dependent most strongly on scan resolution. Biopolymer structures were affected by hydration, as seen in the (c) dry state and (d) hydrated state; (1) slice of porous structure from a μCT scan (< 5 μm voxel size) and (2) 3D reconstruction of the porous features. Significant changes were noted with hydration in (e) wall thickness and (f) percent of the structure occupied by the matrix walls. * p < 0.05, ** p < 0.01, *** p < 0.001. Scale: (a) 1 mm, (c-d) 200 μm.

Since the contrast between polymers and their background disappears once hydrated, treatment with radiopaque contrast agents is necessary to visualize the matrix. Some choose to use liquid contrast agents that can intercalate into the polymer structure [25]. However, this is not necessarily viable when attempting to image polymer devices over a longer period of time or post-implantation. Radiopaque nanoparticles offer a facile way to introduce a radiopaque element into the matrix, without significantly altering materials properties or its biocompatibility [13,26]. They have been shown to work at the scale of clinical tomography and aide in the radiological evaluation of biomedical implants [9].

A caveat to the use of nanoparticle contrast agents is the requirement to demonstrate a truly homogeneous dispersion within a polymer matrix. Otherwise, at the resolution required for visualizing biopolymers, the results could contain too much noise to distinguish features with confidence. Accurate high-resolution scans are capable of capturing changes in small features like swelling of struts in a biopolymer scaffold, Figure 4(c-f). Strut features are highly affected by hydration, with mean thickness increasing from 11.09 ± 3.3 to 26.14 ± 2.9 μm, a 130% increase. The change correlates to a significant reduction in scaffold porosity between the dry and hydrated state, from 92.6 ± 2 to 83.1 ± 4%, respectively. In literature, second harmonic two-photon microscopy has been used to look at the changes in structure after hydration, including an increase in pore size of 20% [23]. This study saw an average of 12% increase in pore size with hydration, in line with literature values. As a caveat, given the high resolution required to visualize swelling of the biopolymer matrix post-hydration, this is a feature that can only be studied in vitro, as it is outside of current resolution of pre-clinical systems on living tissue.

In comparison, the synthetic polymer showed no changes with hydration, Figure 4(e-f). Again, resolution of the scan was a determining factor for the measured pore size and porosity, Figure 4(b). While nothing changed overall, there was a tendency for the variation of measured features to increase in the hydrated state at larger voxel sizes.

Resolving features in biomedical devices at the pre-clinical and clinical scale is not trivial. The challenge of tracking changes in porous structures after implantation necessitates incorporation of contrast agents. If built into the device itself, radiopaque contrast can enable serial monitoring throughout the process of clinical translation. The use of serial monitoring has already been shown to allow tracking of degradation kinetics and failure mode [9,13]. Lessons learned in this space can also be extrapolated to other engineering systems incorporating porosity, including catalysts and battery components [27]. The positive impact of imaging via tomography on biomedical device design could facilitate personalized medicine through individualized rehabilitation and potential intervention post failure.

## 4. CONCLUSIONS

Computed tomography offers a unique tool for biomedical researchers, in that it can provide non-destructive, 3D quantification of porous structures with the ability to reach sub-micron resolutions and can be used to probe features from the nano to macro scale, depending on the tomography system used. Practically speaking, in the medical field, the attainable resolution from CT decreases as the technology moves from pre-clinical to clinical systems, altering the features that can be tracked in porous devices. The poor x-ray attenuation of polymers, often used to create biomedical devices, creates challenges in optimizing scanning parameters for maximum accuracy. In pre-clinical CT scanners, pore size and porosity is more heavily dependent on resolution (voxel size) as opposed to scan time (step size). Another challenge is imaging polymeric devices post-implantation, or in hydrated states, as they are indistinguishable from tissue. The homogeneous introduction of radiopaque TaOx nanoparticles into biopolymers and synthetic polymers allowed tracking changes to porous features during hydration, which are significant in the case of biopolymer matrices. By addressing the issues of resolution and in situ contrast, CT is an imaging technique that is available for monitoring devices through the entire clinical translation pipeline, from the benchtop to the bedside.

## ACKNOWLEDGMENT

The authors would like to thank J. M. L. Hix for help with the in situ implantation models and the Center for Advanced Microscopy (Michigan State University) for help obtaining electron microscopy images. This study was supported by the National Institute of Biomedical Imaging and Bioengineering of the NIH under award number R01EB029418. The content is solely the responsibility of the authors and does not necessarily represent the official views of the National Institutes of Health.

## REFERENCES

[1] Pawelec, K. M.; Hix, J. M. L.; Shapiro, E. M.; Sakamoto, J. The mechanics of scaling-up multichannel scaffold technology for clinical nerve repair. J. Mech. Behav. Biomed. Mater. 2019, 91, 247–254.

[2] Pawelec, K. M.; van Boxtel, H. A.; Kluijtmans, S. G. J. M. Ice-templating of anisotropic structures with high permeability. Mater. Sci. Eng. C. 2017, 76, 628–636.

[3] Sneha, K. R.; Sailaja, G. S. Intrinsically radiopaque biomaterial assortments: a short review on the physical principles, x-ray imageability, and state-of-the-art developments. J. Mater. Chem. B 2021, 9, 8569.

[4] Hemmatian, H.; Laurent, M. R.; Ghazanfari, S.; Vandershueren, D.; Bakker, A. D.; Klein-Nulend, J.; van Lenthe, G. H. Accuracy and reproducibility of mouse cortical bone microporosity as quantified by desktop microcomputed tomography. PloS ONE 2017,12, e0182996.

[5] Wallyn, J.; Anton, N.; Akram, S.; Vandamme, T. F. Biomedical imaging: principles, technologies, clinical aspects, contrast agents, limitations and future trends in nanomedicines. Pharm. Res. 2019, 36, 78.

[6] Tabor, Z. Analysis of the influence of image resolution on the discriminating power of trabecular bone architectural parameters. Bone 2004, 34, 170–179.

[7] Cengiz, I. F.; Oliveria, J. M.; Reis, R. L. Micro-computed tomography characterization of tissue engineering scaffolds: effects of pixel size and rotation step. J. Mater. Sci: Mater. Med. 2017, 28, 129.

[8] Morris, D. E.; Mather, M. L.; Simon, C. G.; Crow, J. A. Time-optimized x-ray microCT imaging of polymer based scaffolds. J. Biomed. Mater. Res. Part B 2012, 100B, 360–367.

[9] Pawelec, K. M.; Chakravarthy, S.; Hix, J. M. L.; Perry, K.; van Holsbeeck, L.; Fajardo, R.; Shapiro, E. Design consideration to facilitate clinical radiological evaluation of implantable biomedical structures. ACS Biomater. Sci. Eng. 2021, 7(2), 718–726.

[10] Veronesi, F.; Di Matteo, B.; Vitale, N. D.; Filardo, G.; Visani, A.; Kon, E.; Fini, M. Biosynthetic scaffolds for partial meniscal loss: A systematic review from animal models to clinical practice. Bioactive Mater. 2021, 6, 3782–3800.

[11] Pawelec, K. M.; Koffler, J.; Shahriari, D.; Galvan, A.; Tuszynski, M. H.; Sakamoto, J. Microstructure and in vivo characterization of multi-channel nerve guidance scaffolds. Biomed. Mater. 2018, 13, 044104.

[12] Chakravarty, S.; Hix, J. M. L.; Wiewiora, K. A.; Volk, M. C.; Kenyon, E.; Shuboni-Mulligan, D. D.; Blanco-Fernandez, B.; Kiupel, M.; Thomas, J.; Sempere, L. F.; Shapiro, E. M. Tantalum oxide nanoparticles as versatile contrast agents for x-ray computed tomography. Nanoscale 2020, 12, 7720–7734.

[13] Pawelec, K. M.; Tu, E.; Chakravarty, S.; Hix, J. M. L.; Buchanan, L.; Kenney, L.; Buchanan, F.; Chatterjee, N.; Das, S.; Alessio, A.; Shapiro, E. M. Incorporating tantalum oxide nanoparticles into implantable polymeric biomedical devices for radiological monitoring. Adv. Healthcare Mater. 2023, 12(18), 2203167.

[14] Bartnikowski, M.; Dargaville, T. R.; Ivanovski, S.; Hutmacher, D. W. Degradation mechanisms of polycaprolactone in the context of chemistry, geometry and environment. Prog. Polym. Sci. 2019, 96, 1–20.

[15] Pawelec, K. M.; Husmann, A.; Best, S. M.; Cameron, R. E. Understanding anisotropy and architecture in ice-templated biopolymer scaffolds. Mater. Sci. Eng. C 2014, 37, 141–147.

[16] Pawelec, K. M.; Husmann, A.; Best, S. M.; Cameron, R. E. A design protocol for tailoring ice templated scaffold structure. J. Royal. Soc. Interface. 2014, 11, 20130958.

[17] Palacio-Mancheno, P. E.; Lariera, A. I.; Doty, S. B.; Cardoso, L.; Fritton, S. P. 3D assessment of cortical bone porosity and tissue mineral density using high-resolution μCT: Effects of resolution and threshold method. J. Bone Mineral Res. 2014, 29(1), 142–150.

[18] Shao, Y.-H.; Tsai, K.; Kim, S.; Wu, Y.-J.; Demissie, K. Exposure to tomographic scans and cancer risks. JNCI Cancer Spectrum 2020, 4(1), pkz072.

[19] Meganck, J.; Liu, B. Dosimetry in micro-computed tomography: a review of the measurement methods, impacts, and characterization of the Quantum GX. Mol. Imaging Biol. 2017, 19, 499–511.

[20] Kerckhofs, G.; Durand, M.; Vangoitsenhoven, R.; Marin, C.; van der Schueren, B.; Caremliet, G.; Luyten, F. P.; Geris, L.; Vandamme, K. Changes in bone macro- and microstructure in diabetic obese mice revealed by high resolution microfocus x-ray computed tomography. Sci. Rep. 2016, 6, 35517.

[21] Alberich-Bayarri, A.; Moratal, D.; Escobar Ivirico, J. L.; Rodriguez Hernandez, J. C.; Valles-Lluch, A.; Marti-Bonmati, L.; Mas Estelles, J.; Mano, J. F.; Monleon Pradas, M.; Gomez Ribelles, J. L.; Salmeron-Sanchez, M. Microcomputed tomography and mirofinite element modeling for evaluating polymer scaffolds architecture and their mechanical properties. J. Biomed. Mater. Res Part B 2009, 91B, 191–202.

[22] Pawelec, K. M.; Husmann, A.; Best, S. M.; Cameron, R. E. Ice-templated structures for biomedical tissue repair: From physics to final scaffolds. Appl. Phys. Rev. 2014, 1, 021301.

[23] Varley, M.; Neelakantan, S.; Clyne, T. W.; Dean, J.; Brooks, R. A.; Markaki, A. E. Cell structure, stiffness and permeability of freeze-dried collagen scaffolds in dry and hydrated states. Acta Biomater. 2016, 33, 166–175.

[24] Fager, C.; Barman, S.; Roeding, M.; Olsson, A.; Loren, N.; von Corswant, C.; Bolin, D.; Rootzen, H.; Olsson, E. 3D high spatial resolution visualization and quantification of interconnectivity in polymer films. Int. J. Pharm. 2020, 587, 119622.

[25] Suchy, T.; Supova, M.; Bartos, M.; Sedlacek, R.; Piola, M.; Soncini, M.; Fiore, G. B.; Sauerova, P.; Kalbacova, M. H. Dry versus hydrated collagen scaffolds: are dry states representative of hydrated states? J. Mater. Sci: Mater. Med. 2018, 29, 20.

[26] Pawelec, K. M.; Hix, J. M. L.; Shapiro, E. M. Functional attachment of primary neurons and glia on radiopaque implantable biomaterials for nerve repair. Nanomed.: Nanotechnol. Biol. Med. 2023, 52, 102692.

[27] Scharf, J.; Chouchane, M.; Finegan, D. P.; Lu, B.; Request, C.; Kim, W.; Yao, M-C.; Franco, A.; Gostovic, D.; Liu, Z.; Riccio, M.; Zelenka, F.; Doux, J.-M.; Meng, Y. S. Bridging nano- and microscale x-ray tomography for battery research by leveraging artificial intelligence. Nat. Nanotechnol. 2022, 17, 446–459.

